# IL-33-induced neutrophilic inflammation and NETosis underlie rhinovirus-triggered exacerbations of asthma

**DOI:** 10.1101/2022.11.07.515526

**Authors:** Bodie Curren, Tufael Ahmed, Daniel R Howard, Md. Ashik Ullah, Ismail Sebina, Ridwan B Rashid, Md. Al Amin Sikder, Alec Bissell, Sylvia Ngo, David J Jackson, Marie Toussaint, Michael R. Edwards, Sebastian L Johnston, Henry J. McSorley, Simon Phipps

**Author notes:** Joint First Authors (Bodie Curren, Tufael Ahmed). Correspondence: Dr. Simon Phipps, QIMR Berghofer Medical Research Institute, Herston, Queensland, 4006, Australia.

## Abstract

Rhinovirus-induced neutrophil extracellular traps (NETs) contribute to acute asthma exacerbations, however the molecular factors that trigger NETosis in this context remain ill-defined. Here, we sought to implicate a role for IL-33, an epithelial cell-derived alarmin rapidly released in response to infection. In mice with chronic experimental asthma (CEA), but not naïve controls, rhinovirus inoculation induced an early (1 day post infection; dpi) inflammatory response dominated by neutrophils, neutrophil-associated cytokines (IL-1α, IL-1β, CXCL1) and NETosis, followed by a later, type-2 inflammatory phase (3-7 dpi), characterized by eosinophils, elevated IL-4 levels, and goblet cell hyperplasia. Notably, both phases were ablated by HpARI (*Heligmosomoides polygyrus* Alarmin Release Inhibitor), which blocks IL-33 release and signalling. Instillation of exogenous IL-33 recapitulated the rhinovirus-induced early phase, including the increased presence of NETs in the airway mucosa, in a PAD4-dependent manner. *Ex vivo* IL-33-stimulated neutrophils from mice with CEA, but not naïve mice, underwent NETosis, and produced greater amounts of IL-1α/β, IL-4, and IL-5. In nasal samples from rhinovirus-infected people with asthma, but not healthy controls, IL-33 levels correlated with neutrophil elastase and dsDNA. Our findings suggest that IL-33 blockade ameliorates the severity of an asthma exacerbation by attenuating neutrophil recruitment and the downstream generation of NETs.

## Introduction

Asthma is a chronic respiratory disease that affects 400 million people worldwide, and is typically underpinned by the generation of aberrant type-2 immune responses.^1^ Acute exacerbations of asthma, which cause significant morbidity and associated health care costs, are commonly triggered by a respiratory virus, mostly rhinovirus (RV) infection, amplifying pathogenic type-2 inflammation. Amongst the earliest events of a viral-triggered exacerbation is the increased expression of neutrophil-active chemokines, such as CXCL8, and correspondingly, an influx of neutrophils to the lungs and airways.^2, 3^ We found that this neutrophilic response, through the release of neutrophil extracellular traps (NETs), promotes the pathogenic type-2 inflammation that induces the cardinal features of asthma.^4^ Significantly, the therapeutic targeting of NETs, extruded nuclear DNA bound with neutrophil elastase and Citrullinated histone-H3 (Cit-H3),^5^ via the administration of DNase ameliorated the severity of RV-induced exacerbation in a mouse model of experimental asthma.^4^ Although this study identified NETosis as a critical intermediary of RV-induced type-2 inflammation and ensuing loss of asthma control, the events in the airway mucosa that promote NETosis in the context of asthma remain poorly defined.

In subjects with asthma, mucosal homeostasis is dysregulated. This is typified by an airway epithelium that is hyperplastic, contains greater numbers of mucus-secreting goblet cells, and sits on a thickened reticular basement membrane.^6^ Additionally, the epithelium initiates and shapes the nature of the immune response through the production of inflammatory mediators, such as eicosanoids, chemokines, and type-2-instructive cytokines such as thymic stromal lymphopoietin (TSLP) and interleukin-33 (IL-33). The targeting of these epithelial-derived alarmins with monoclonal antibodies in clinical trials has been shown to dampen airway inflammation, improve lung function, and decrease the frequency of exacerbations,^7, 8^ highlighting the value of fundamental insights into type-2 immunity learnt from clinical and experimental model systems.^9–11^ However, although the efficacy of these biologics is particularly potent in individuals with eosinophilic or allergic phenotypes,^12^ a recent Phase II trial found that targeting ST2, one of the IL-33 receptor chains, significantly lowers the frequency of exacerbations in individuals with uncontrolled severe type-2-low asthma (i.e. patients with low blood eosinophils).^13^ Additionally, preclinical models of chronic experimental asthma (CEA) show that as well as inhibiting type-2 inflammation, anti-IL-33 treatment attenuates neutrophilic inflammation.^9, 10^ Collectively, these data suggest that IL-33 additionally contributes to non-type-2 inflammation in asthma, although whether IL-33 promotes neutrophil recruitment and NETosis in asthma remains unknown.

We previously developed and characterised a unique mouse model of CEA that simulates the human epidemiology linking severe lower respiratory infections in early life to the later development of allergic asthma.^9, 14, 15^ Through repeated virus and allergen exposure in early and later life, the mice develop CEA, as demonstrated by fixed airway remodelling and persistently high levels of airway IL-33, and significantly, are predisposed to a RV-induced exacerbation characterized by an early neutrophilia that peaks at 24 hours, followed by a type-2 inflammatory response and mucus hypersecretion that increases over days.^9, 15^ Using this model and HpARI (*Heligmosomoides polygyrus* Alarmin Release Inhibitor), a secretory worm product that we have previously shown to inhibit both IL-33 release and downstream ST2 signalling,^16^ we sought to test the hypothesis that in the setting of CEA, RV-induced IL-33 release amplifies type-2 inflammation via the recruitment of neutrophils and the induction of NETosis.

## Results

### HpARI suppresses RV-induced acute exacerbation of chronic experimental asthma

We previously demonstrated that repeated exposures to the mouse respiratory virus, pneumonia virus of mice (PVM; related to human respiratory syncytial virus), and cockroach allergen in early and later life, simulating the human epidemiology,^17^ leads to the onset and development of chronic experimental asthma (CEA) with persistent airway remodelling.^14^ Importantly, in mice with established asthma, but not naïve controls, inoculation with RV induces an inflammatory response that is characteristic of an acute exacerbation of asthma.^9, 15^ This model of CEA is characterized by persistently elevated IL-33 expression in the airways that increases further in response to RV infection together with an early neutrophilic response,^9^ leading us to explore whether IL-33 blockade through intranasal administration of HpARI would ameliorate both the RV-induced neutrophilia and the ensuing type-2 inflammation (Fig. 1A). Confirming and extending our previous observations, mice with CEA, but not age-matched naive controls, developed a significant goblet cell hyperplasia and a mixed type-2/type-17 inflammatory response post inoculation with RV (Fig. 1B-E and Supplementary Fig. 1). Notably, neutrophils were the predominant granulocyte at 1 dpi, outnumbering eosinophils 8 to 1 (Fig.1C). Siglec-F+ neutrophils, reported by some investigators, were rare to absent (data not shown). Following HpARI-treatment, this pronounced, early wave of infiltrating neutrophils to the lungs was markedly attenuated, as was the magnitude of the neutrophil and eosinophil response across the time course (Fig. 1B-C). This was associated with a significant decrease in the numbers of type-2 innate lymphoid cells (ILC2s), CD4+ Th2, and CD4+ Th17 cells, a concomitant decrease in IL-4, IL-5, IL-13 and IL-17A concentrations in bronchoalveolar lavage (BAL) fluid and mucus production (Fig. 1B, D-E). In contrast to ILC2s, ILC3 numbers in the lung were low and unaffected by HpARI (Fig. 1D).

**Figure 1:**
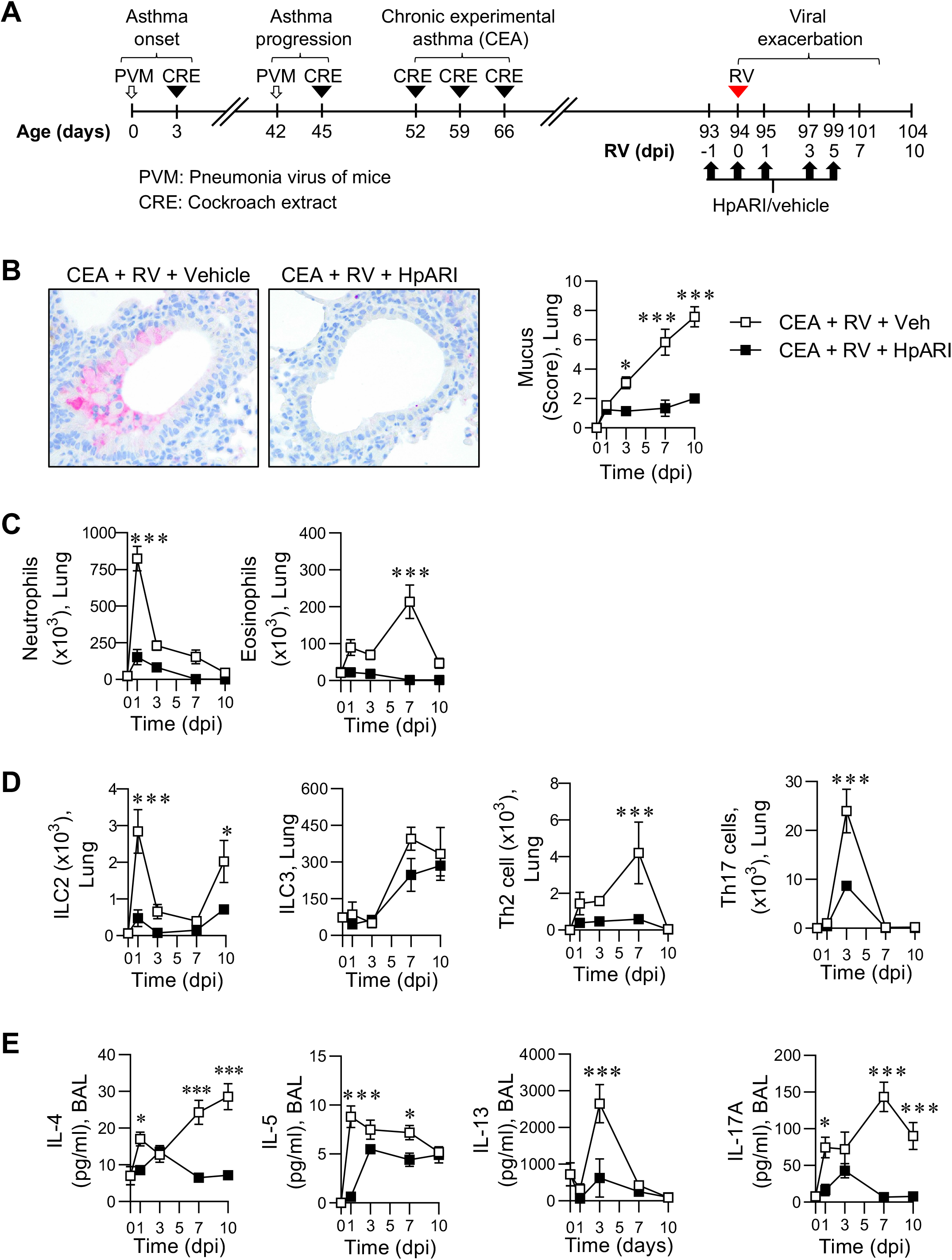
HpARI suppresses an RV-induced acute exacerbation of CEA. **A,** Study design (dpi, day post infection). **B,** Representative images of MUC5ac immunohistochemistry (x20 magnification). MUC5ac score. **C,** Lung neutrophils and eosinophils. **D,** ILC2s, ILC3s, Th2 cells and Th17 cells in the lung. **E,** IL-4, IL-5, IL-13 and IL-17A expression in BAL fluid. Data represented as mean ± SEM, *n* = 4-12 mice per group, \**P* < .05, *** *P* < .00.1.

### HpARI decreases the expression of IL-33 and other innate instructive cytokines in the airways

We next examined the effect of HpARI treatment on the levels of IL-33 and other innate type-2/17 associated-instructive cytokines. Consistent with our previous findings,^9^ IL-33 was highly expressed in mice with CEA prior to RV inoculation (Fig. 2A). Following treatment with HpARI, IL-33 levels in the airway were ablated, an effect that persisted until 10 dpi, 5 days after the final HpARI administration. HpARI also decreased the levels of other instructive cytokines and chemokines, including TSLP, IL-1α, IL-1ß, IL-6 and CXCL1 (Fig. 2B-F), consistent with the attenuated production of type-2/type-17 effector cytokines and ablated granulocytic response in HpARI treated mice, indicating that IL-33 is central to the early inflammatory response.

**Figure 2:**
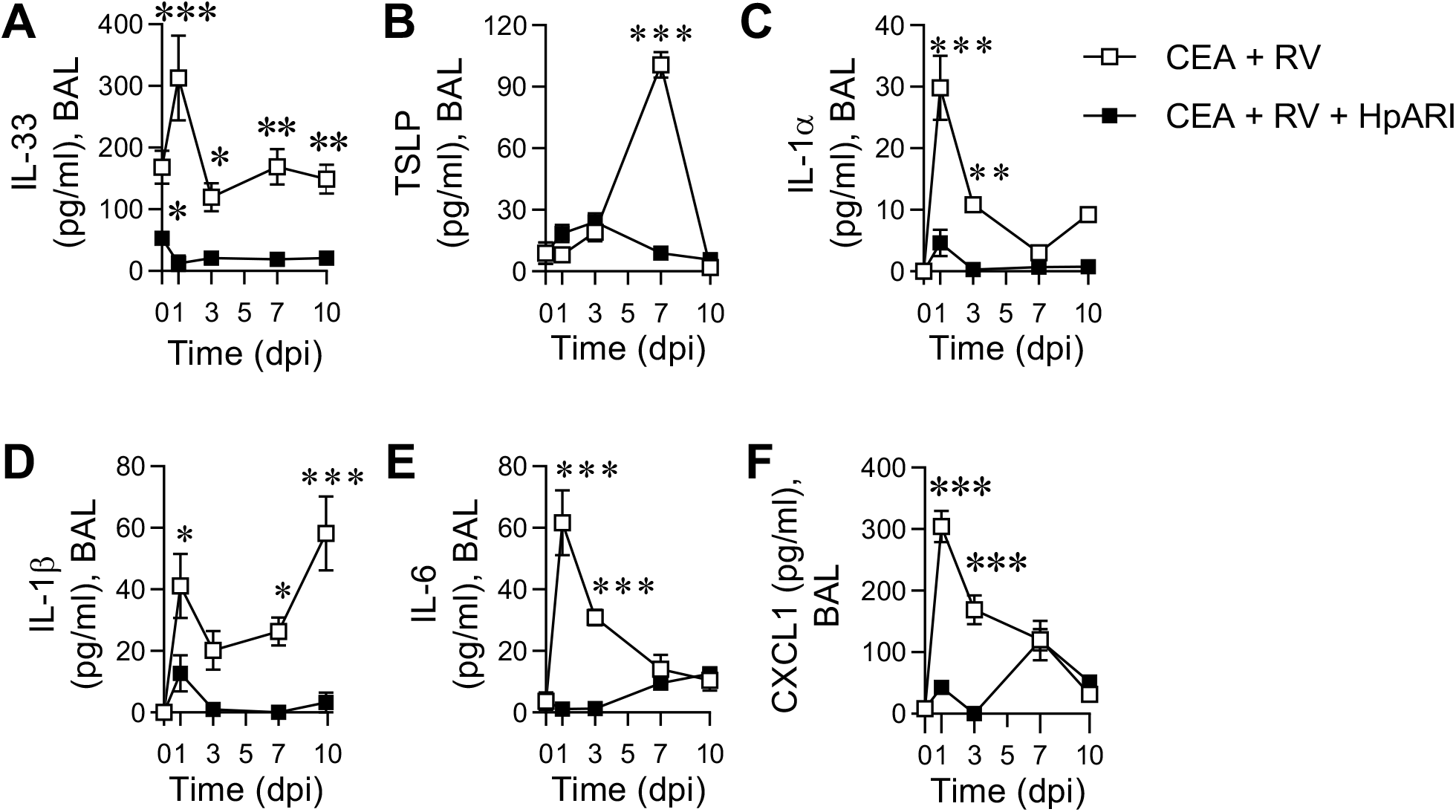
HpARI decreases the expression of IL-33 and other innate instructive cytokines in the airways. **A,** IL-33, **B,** TSLP, **C,** IL-1α, **D,** IL-1β, **E,** IL-6 and **F,** CXCL1 expression in BAL fluid. Data represented as mean ± SEM, *n* = 4-8 mice per group, \**P* < .05, \*\**P* < .01, *** *P* < .00.1.

### Exogenous IL-33 recapitulates the response to RV inoculation in mice with CEA

As HpARI decreased both RV-induced neutrophilic and eosinophilic inflammation, we next assessed whether exogenous IL-33 was sufficient to induce a lung neutrophilia (Fig. 3A). In contrast to naïve mice, where IL-33 induced a mild neutrophilic response and minimally affected lung eosinophil numbers, IL-33 inoculation of mice with CEA triggered a marked neutrophilic response, similar to that observed following RV exposure (Fig. 3A). This response was associated with increased numbers of ILC2s, ILC3s, and CD4+ Th2 cells in the lung, increased production of type-2/17 cytokines and goblet cell hyperplasia, whereas in naïve mice, exogenous IL-33 elicited a limited inflammatory response (Fig. 3B-E). Of note, similar to RV inoculation, exogenous IL-33 increased the concentrations of TSLP, IL-1α, IL-1β, IL-6 and CXCL1 in the BAL fluid (Fig. 4). Collectively, these findings demonstrate that exogenous IL-33 induces a similar response to that elicited by RV inoculation.

**Figure 3:**
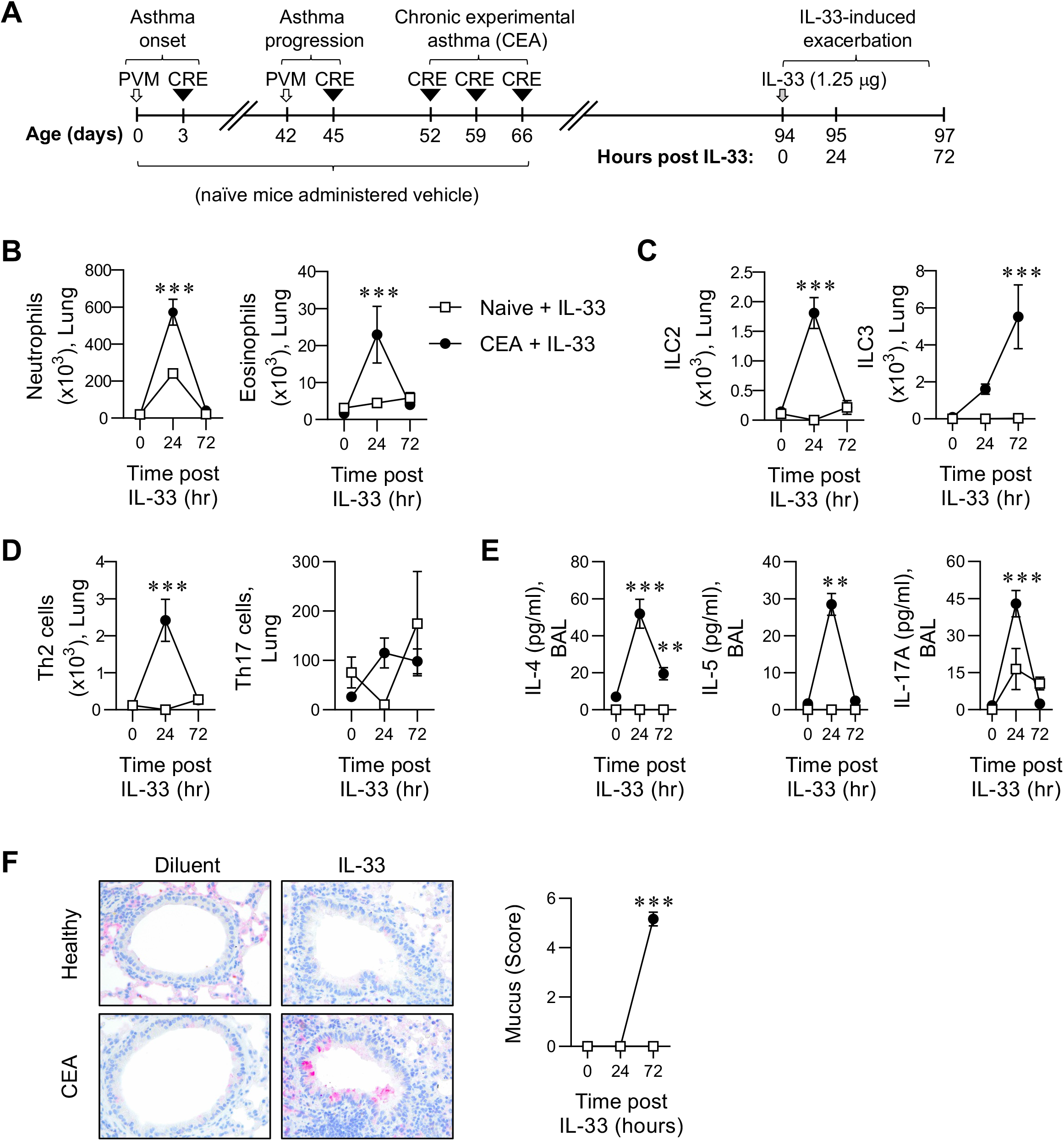
Exogenous IL-33 recapitulates the response to RV inoculation in mice with CEA. **A,** Study design. At postnatal day 7, WT BALB/C mice were inoculated with PVM (1 pfu) or diluent at day 0 then inoculated with CRE (1 μg) or diluent at day 3. Mice were re-infected with PVM (20 pfu) at day 42, and challenged with CRE (1 μg) at day 45, 52, 59 and 66. Naïve mice were inoculated with diluent at the aforementioned time points. At day 94, mice were inoculated with IL-33 (1.25 μg). **B,** Lung neutrophils and eosinophils. **C,** Lung ILC2s and ILC3s. **D,** Lung Th2 cells and Th17 cells. **E,** IL-4, IL-5, and IL-17A expression in BAL fluid. **F,** MUC5ac score. Data represented as mean ± SEM, *n* = 4-8 mice per group, \*\**P* < .01, *** *P* < .00.1.

**Figure 4:**
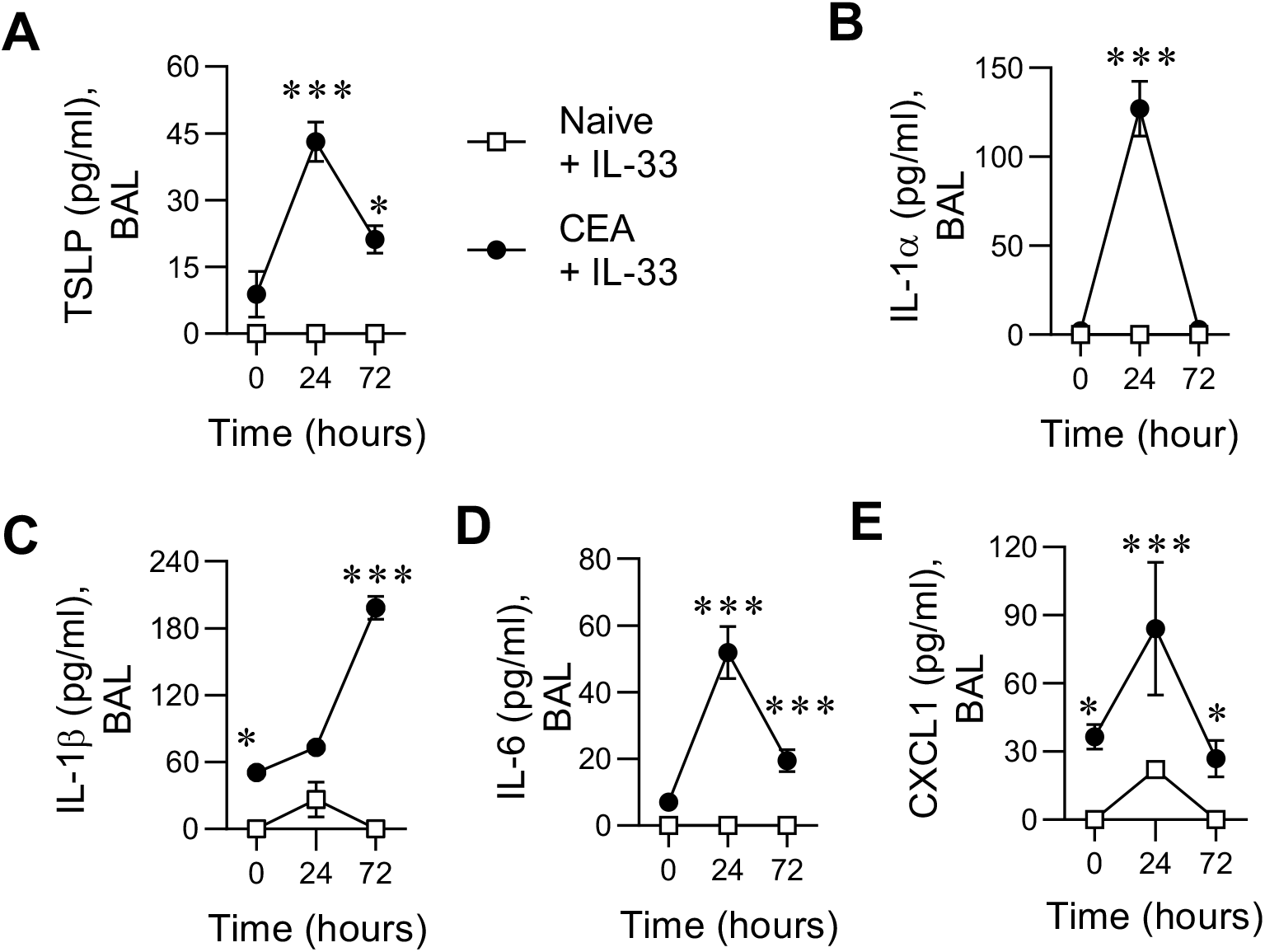
IL-33 promotes the expression of innate instructive cytokines in mice with CEA but not naïve mice. **A,** TSLP, **B,** IL-1α, **C,** IL-1β, **D,** IL-6 and **E,** CXCL1 expression in BAL fluid. Data represented as mean ± SEM, *n* = 4-8 mice per group, \**P* < .05, *** *P* < .00.1.

### RV or IL-33 induced NETosis is attenuated by HpARI

We previously linked NETosis to the amplification of type-2 inflammation.^4^ To assess the effect of HpARI on RV-induced NET formation in the lungs, we performed double immunohistochemistry for myeloperoxidase (MPO) and Citrullinated histone-H3 (CitH3), and quantified the co-localised signal. As expected, RV inoculation of mice with CEA led to a marked increase in co-localised MPO/CitH3 expression, and this response was attenuated in mice treated with HpARI (Fig. 5A). Significantly, exogenous IL-33 also increased MPO/CitH3 co-localisation, indicative of NETosis, but only in mice with CEA (Fig. 5B). Additionally, dsDNA levels were elevated in the airways in response to RV or IL-33 inoculation in mice with CEA, but not in naïve controls, and the concentration of dsDNA was significantly decreased in RV inoculated CEA mice following treatment with HpARI (Fig. 5C-D), further supporting the notion that IL-33 is a key mediator of NETosis in mice with CEA.

**Figure 5:**
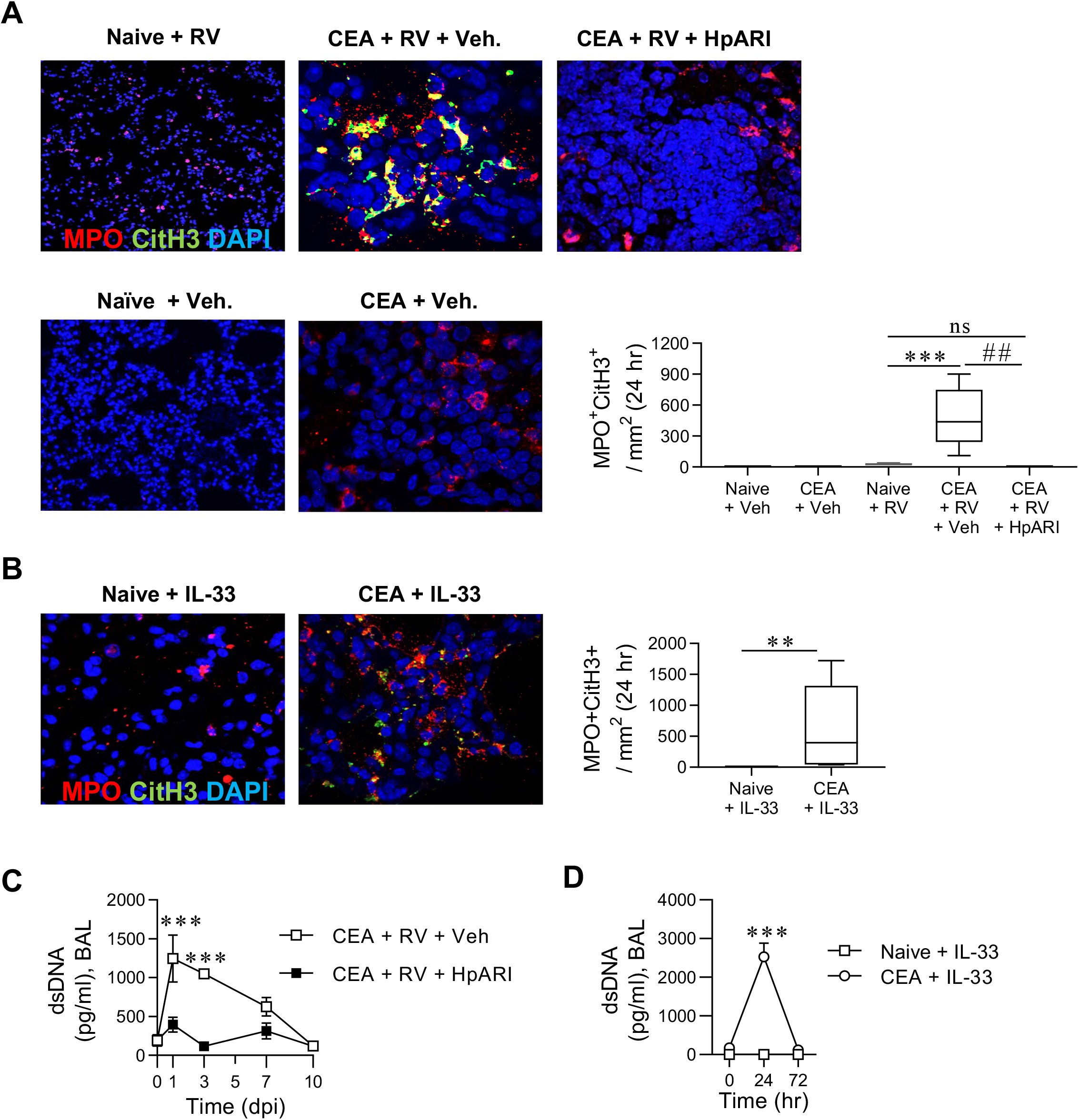
RV or IL-33 inoculation induced NETosis is attenuated by HpARI. **A,** Representative immunofluorescent images of myeloperoxidase (MPO; red), Citrullinated histone-H3 (CitH3: green), and DNA (DAPI; blue). Quantification of co-localised MPO and CitH3 expression in lung tissue at 1 dpi (day post infection). **B,** Representative immunofluorescent images of MPO (red), CitH3 (green), and DNA (DAPI; blue). Quantification of co-localised MPO and CitH3 expression in lung tissue at 1 day post IL-33 exposure. **C,** dsDNA levels in BAL fluid of CEA mice following RV infection ± HpARI. **D,** dsDNA levels in BAL of CEA or naïve mice following IL-33 exposure. Data represented as mean ± SEM, *n* = 4-8 mice per group, *\*\*P* < .01, *** *p* < .00.1 (compared to Naïve + Veh), ## *P* < .01 (compared to CEA + RV + Veh).

### IL-33 associates with dsDNA and neutrophil elastase following experimental RV challenge of people with mild to moderate asthma

We previously demonstrated that the concentrations of dsDNA and neutrophil elastase, markers of NETosis, were significantly greater in the nasal fluid of subjects with asthma compared to healthy controls following experimental RV infection. Here we found that peak nasal IL-33 concentration significantly correlated with peak nasal neutrophil elastase as well as peak nasal dsDNA, two biomarkers of NETosis, following RV infection only in subjects with asthma, but not in healthy control subjects (Fig. 6A-B). Moreover, at day 3 post RV infection, the concentration of nasal dsDNA positively correlated with nasal IL-33 levels (Fig. 6C), as well as upper and lower respiratory symptoms, and again only in subjects with asthma, but not in healthy control subjects (Fig. 6D-E).

**Figure 6:**
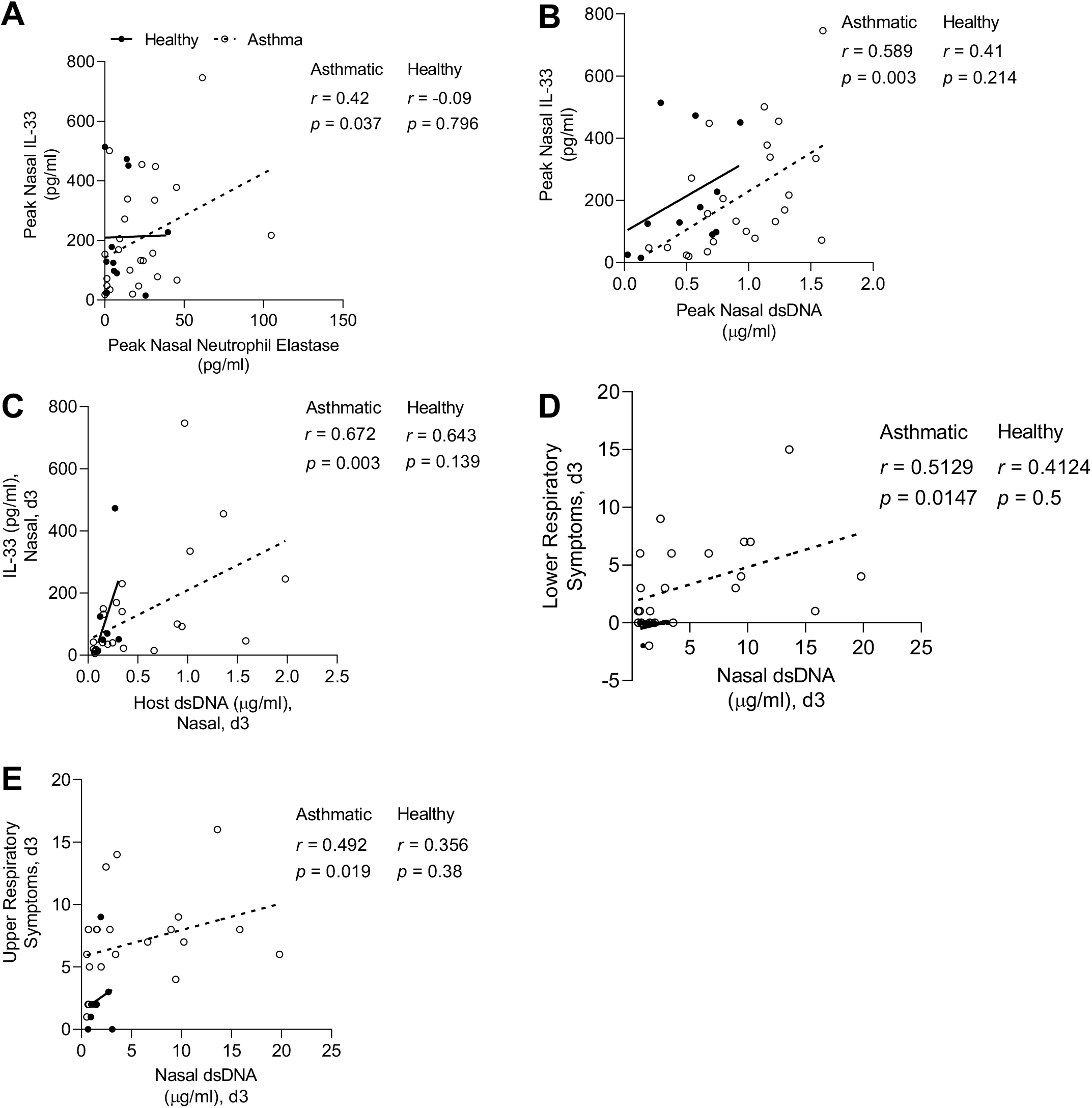
IL-33 associates with dsDNA and neutrophil elastase following experimental RV challenge of mild to moderate asthmatics. Nasal levels of IL-33, neutrophil elastase, host dsDNA, upper respiratory symptoms and lower respiratory symptoms were measured in 24 subjects with asthma and 11 healthy controls. **A,** Correlation between peak levels of nasal IL-33 and peak levels of nasal neutrophil elastase (i.e., maximal levels of cytokines during infection for each subject). **B,** Correlation between peak nasal IL-33 and peak nasal host dsDNA. **C,** Correlation between nasal concentrations (on day 3 during RV infection) of IL-33 and host dsDNA. **D,** Correlation between lower respiratory symptoms (on day 3 during RV infection) and nasal host dsDNA. **E,** Correlation between upper respiratory symptoms (on day 3 during RV infection) and nasal host dsDNA. **A-E,** The correlation analysis used was nonparametric (Spearman’s correlation) performed on subjects with asthma and healthy controls. \**P* < .05, \*\**P* < .01, *** *P* < .00.1.

### NETosis inhibition attenuates an IL-33-induced exacerbation of chronic experimental asthma

Since exogenous IL-33 induced NETosis, we hypothesized that inhibition of NETosis using a Protein arginine deiminase 4 (PAD4) inhibitor (Supplementary Fig. 2A) would attenuate IL-33-induced type-2/17 inflammation. As expected, PAD4 inhibition significantly decreased the presence of co-localised MPO/CitH3 in the lung tissue and dsDNA levels in the airways (Fig. 7A-B). Consistent with our hypothesis, PAD4 inhibition ablated IL-33-induced granulocytic inflammation, ILC2 and ILC3 numbers, Th2 and Th17 cell numbers, type-2/17 effector cytokines and mucus hyperproduction (Fig. 7C-E and Supplementary Fig. 2B-C). Strikingly, in addition to suppressing type-2/17 inflammation, inhibition of NETosis ablated IL-33-induced IL-1α, IL-1ß, IL-6, and CXCL1 levels in the BAL fluid. In contrast, TSLP levels were unaffected (Fig. 7F).

**Figure 7:**
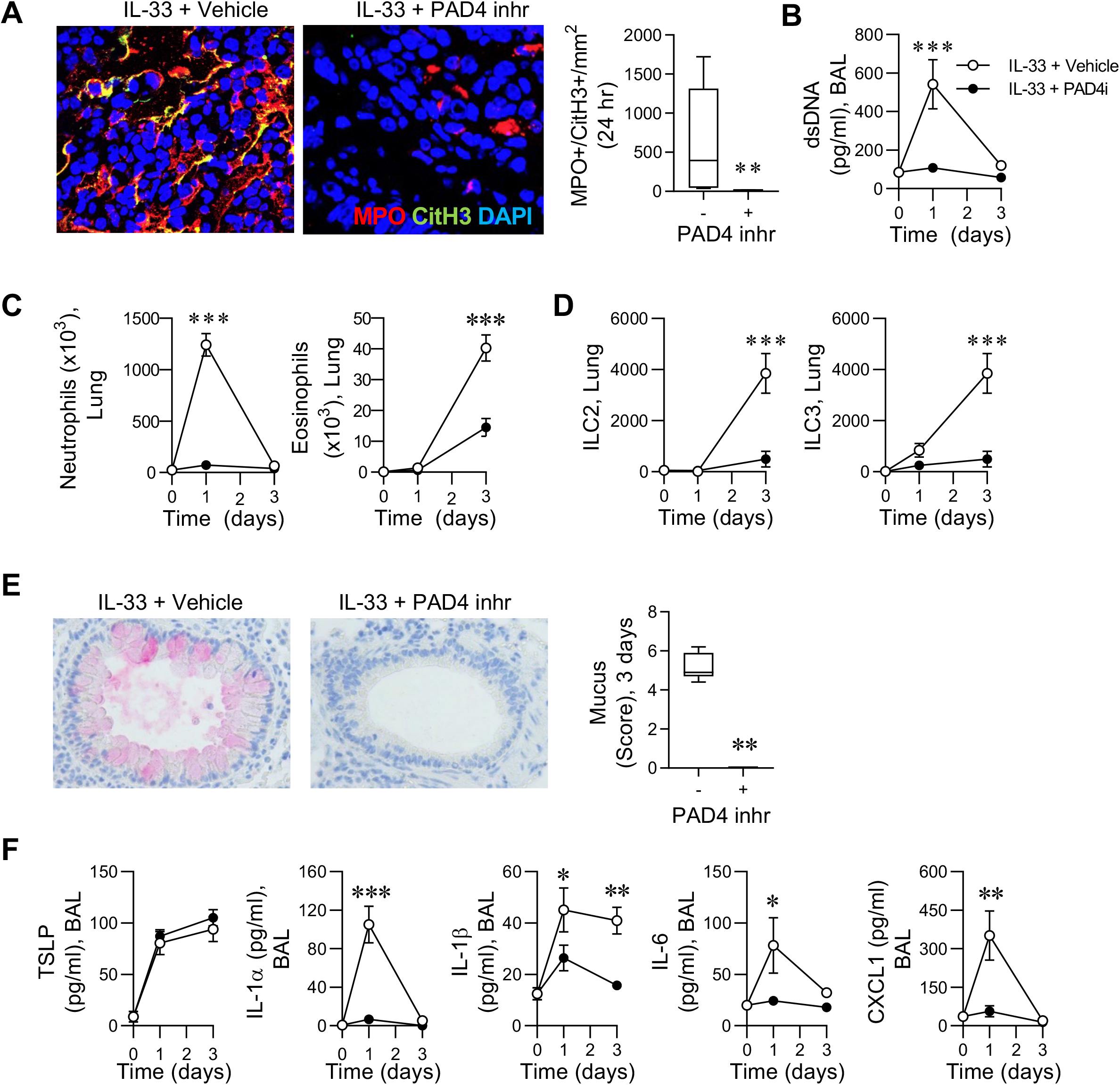
NETosis inhibition attenuates an IL-33-induced exacerbation of CEA. **A,** Representative immunofluorescent images of MPO (red), CitH3 (green), and DNA (DAPI, blue) (x20 magnification). Quantification of co-localised MPO and CitH3 expression in lung tissue 24 hours post IL-33 exposure. **B,** dsDNA levels in BAL. **C,** Lung neutrophils and eosinophils. **D,** Lung ILC2s and ILC3s. **E,** Representative images of MUC5ac IHC (x20 magnification). Mucus score. **F,** TSLP, IL-1α, IL-1β, IL-6 and CXCL1 expression in BAL. Data represented as mean ± SEM, *n* = 4-8 mice per group, *\*P* < .05, *\*\*P* < .01, *** *P* < .00.1.

### IL-33 induces NETosis and asthma related cytokines from neutrophils of mice with CEA but not in naive controls

To test whether IL-33 acts directly on neutrophils to induce NETosis, naïve mice and mice with CEA were inoculated with RV, and then 4 hours later, neutrophils were FACS sorted from the circulation or bone marrow (Supplementary Fig. 3A). Strikingly, in peripheral blood (PB) neutrophils obtained from RV-exacerbated CEA, but not from RV-inoculated naïve mice, IL-33 induced dsDNA release and increased the expression of co-localised MPO/CitH3 (Fig. 8A-B). The same phenotype was observed in bone marrow neutrophils (Supplementary Fig. 3B-C). Lastly, as peak neutrophil infiltration coincided with high airway cytokine expression (i.e. at 1 dpi), we assessed whether the *ex vivo* cultured PB neutrophils released IL-1α, IL-1β, IL-4, IL-5, IL-6, IL-17A and KC. Of note, in the absence of stimulation, IL-1α, IL-1β, IL-4, IL-5, and IL-6 secretion, but not IL-17A or KC (data not shown) was significantly greater from neutrophils purified from mice with CEA, compared to healthy controls (Fig. 8C-D). Critically, IL-33 markedly increased the production of these cytokines by PB neutrophils of mice with CEA (Fig. 8C-D). In contrast, PB neutrophils from naïve mice only produced a small increase in IL-4 production in response to IL-33 (Fig. 8C-D). Collectively, and consistent with our *in vivo* observations, these findings demonstrate that IL-33 acts directly on neutrophils to promote NETosis and cytokine release, and importantly, demonstrate that the propensity for neutrophils to undergo this response is markedly affected by disease status.

**Figure 8:**
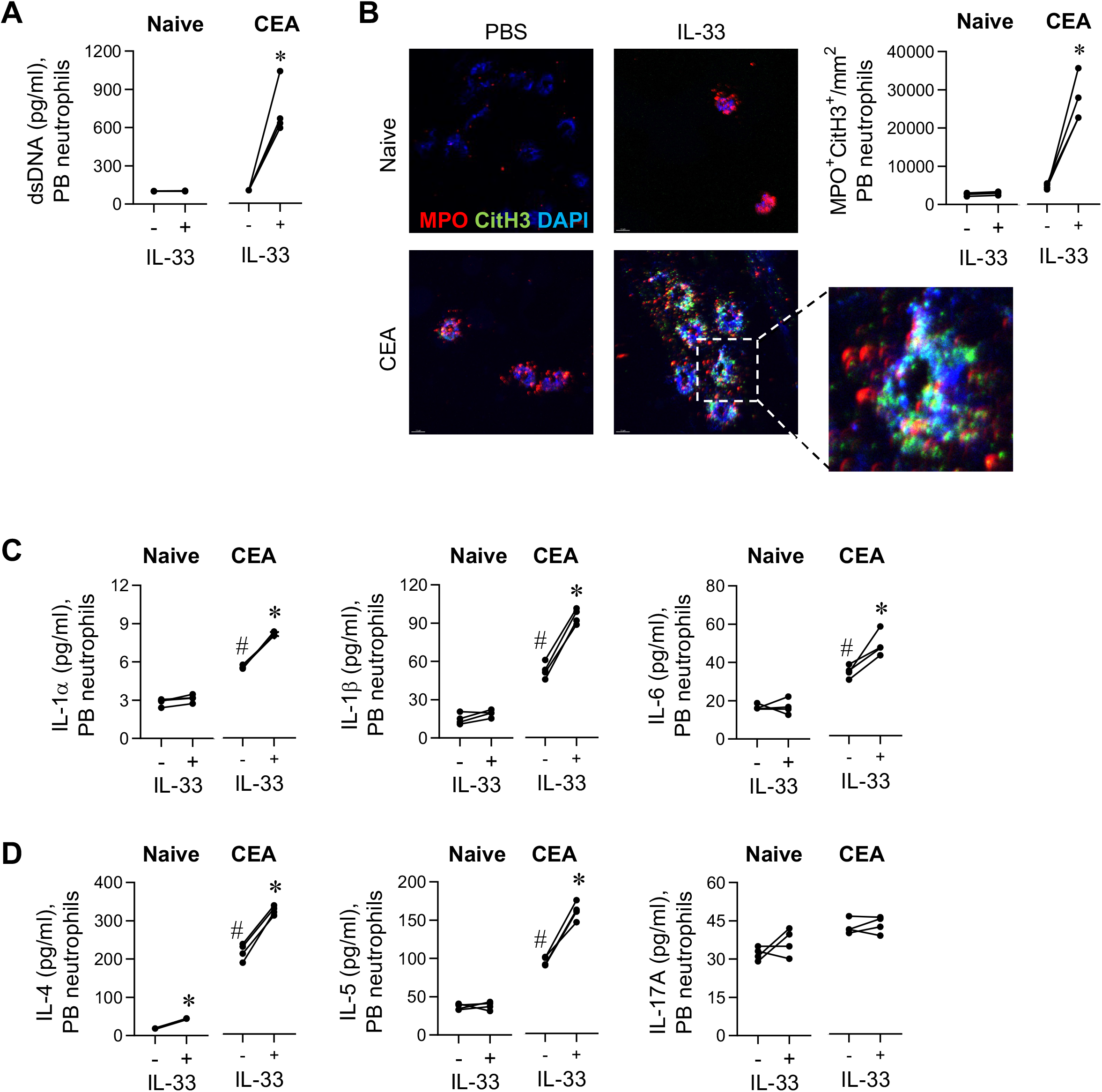
IL-33 induces NETosis and asthmatogenic cytokines from neutrophils of mice with CEA but not healthy controls. **A,** dsDNA levels in the supernatant of peripheral blood (PB) neutrophils isolated from from naïve or CEA mice and stimulated ± IL-33. **B,** Representative images of immunofluorescent MPO (red), CitH3 (green), and DNA (DAPI, blue) in neutrophils stimulated ±IL-33 *ex vivo*. Quantification of co-localised MPO and CitH3 expression. **C**, **D,** IL-4, IL-1α, IL-1β and IL-6 (*C*) and IL-4, IL-5, and IL-17A (*D*) expression in the supernatant of PB neutrophils, stimulated ± IL-33. Data represented as mean ± SEM, *n* = 4-8 mice per group, \**P* < .05, \*\**P* < .01, *** *P* < .00.1 (comparing ± IL-33), ## *P* < .01 (comparing unstimulated naïve to CEA).

## Discussion

Using an established mouse model of CEA that is characterized by a persistent IL-33-high microenvironment in the airways,^9^ we sought to test the efficacy of HpARI, a worm product that neutralizes IL-33 and IL-33 signalling,^16^ during a RV induced exacerbation. As expected, and consistent with our findings with anti-IL-33,^9^ HpARI ablated the expansion of the type-2 inflammation and mucus hyper production that occurs in response to RV inoculation. However, unexpectedly, HpARI ablated the early innate inflammatory response that is dominated by neutrophilic inflammation and associated cytokines, and markedly decreased the presence of NETs in the airway mucosa. Of note, exogenous IL-33 recapitulated the response observed following RV inoculation, including the induction of NETosis by infiltrating neutrophils, and significantly, treatment with a PAD4 inhibitor conferred protection against an IL-33-induced exacerbation. Critically, *ex vivo* IL-33 stimulation induced NETosis and the release of asthma-related cytokines from peripheral blood neutrophils isolated from mice with CEA, whereas neutrophils from naïve mice were unresponsive, suggesting that altered transcriptional and/or chromatin changes are operational prior to neutrophil migration to the lungs. Collectively, our findings identify an important role for IL-33 in promoting neutrophil inflammation and NETosis.

IL-33 is constitutively expressed in the cell nucleus and described as a danger-associated molecular pattern or alarmin as it is released in response to damage, stress, or infection.^19^ The identification of IL-33 as a cytokine that activates effector cells involved in type-2 inflammation (e.g. mast cells, eosinophils, ILC2s) and as an inducer of CD4+ Th2 cell differentiation, together with efficacious findings in response to IL-33 blockade in preclinical mouse models^9, 14, 21^ led to the development of biologics targeting IL-33 and the IL-33 receptor, ST2. Several clinical trials using these agents are on-going, however a recent Phase 2 trial found that anti-IL-33 lowers the incidence of exacerbations and improves lung function in moderate-to-severe asthma,^7^ further supporting a role for IL-33 in type-2 mediated inflammation. Similarly, monoclonal antibody targeting of the ST2 (IL-33) receptor decreased the asthma exacerbation rate in patients with uncontrolled severe asthma. Of interest, this positive outcome was also observed in those with low peripheral blood eosinophils, suggesting that IL-33 blockade may be an effective treatment option for subtypes of asthma less dominated by type-2/eosinophilic inflammation.^13^ Although neither of these clinical studies reported the effect of IL-33 blockade on neutrophil numbers or biomarkers of neutrophil activation, we previously observed that anti-IL-33 attenuates neutrophilic inflammation induced by RV exposure in mice with CEA.^9^ Here, we observed that IL-33 blockade with HpARI ablated the RV-induced inflammatory response which appeared to occur in two phases; the first between 1-3 dpi, and the second between 7-10 dpi. The latter phase, characterized by higher numbers of CD4+ Th2 cells and eosinophils, goblet cell hyperplasia and greater IL-4 expression, was ablated by HpARI, in keeping with the known effects of IL-33 in amplifying type-2 immunity. Significantly, however, treatment with HpARI also suppressed the early neutrophilic response, characterized by a spike in lung neutrophils, innate inflammatory cytokines/chemokines (e.g. IL-1α/β, IL-6, CXCL1), type-2/3 cytokines (e.g. IL-5 and IL-17A), and diminished the number of NETs in the airway mucosa. These findings suggested that IL-33 promotes neutrophilic inflammation, and in support of this, we demonstrated that in mice with CEA, but not naive controls, exogenous IL-33 exposure is sufficient to recapitulate the early phase response induced by RV, including the marked increase in lung neutrophils, NET formation in the mucosa and elevated biomarkers of NETosis in the BAL, and an increase in the expression of inflammatory cytokines/chemokines and type-2/17 cytokines. Importantly, the IL-33-induced inflammatory response and NETosis was ablated with a PAD4 inhibitor, which is significant as NETosis plays a critical pathogenic role in linking RV-induced innate inflammation to the expansion of type-2 immune responses and immunopathology in asthma.^4^ The molecular pathway by which NETosis promotes type 2 inflammation remains unclear, however it is noteworthy that a number of neutrophil-derived factors released during NETosis, such as LL-37, HMGB1, and S100A8/A9, are ligands of receptor for advanced glycation end-products,^22, 23^ an innate receptor that we and others have implicated in the induction and amplification of type 2 immune responses.^24–26^

IL-33 acts directly on human eosinophils to promote their survival and induces the release of reactive oxygen species and cytokines.^27^ In contrast, the effects of IL-33 on human neutrophil function is less clear, with some studies reporting an absence of neutrophil ST2 expression^27^ whereas others have shown that IL-33 regulates CXCR1/CXCR2 expression and CXCL8-mediated chemotaxis, and one report in the context of liver injury has suggested that IL-33 promotes neutrophil NETosis.^28–30^ In support of a direct role for IL-33 in inducing neutrophil activation, we found that *ex vivo* stimulation of purified blood and bone marrow neutrophils was sufficient to induce NETosis but only when the neutrophils were obtained from mice with CEA. In contrast, *ex vivo* IL-33-stimulated neutrophils from naïve mice were unresponsive. This finding was consistent with the *in vivo* phenotype, where intranasal IL-33 exposure induced a marked inflammatory response in mice with CEA but not naïve controls. In subjects experimentally inoculated with RV, nasal dsDNA levels, a biomarker of neutrophil NETosis, correlated with nasal IL-33 levels, as well as respiratory symptoms, in those with asthma, but not healthy controls, further supporting IL-33-induced NETosis as a disease-specific phenomenon. Interestingly, others have similarly reported differences in IL-33 responsiveness between subjects with asthma and healthy controls,^31^ and although the underlying mechanism was not identified, the effect was not mediated by a difference in basal ST2 expression.

In mice with CEA, the large increase in pro-inflammatory cytokine production at 1 dpi occurred concomitantly with peak lung neutrophilia, suggesting that neutrophils may contribute to this phenotype. Remarkably, even in the absence of stimulation, circulating neutrophils obtained from mice with CEA produced significantly greater levels of IL-1β, IL-4, IL-5, and IL-6. Given the enhanced production of type-2 cytokines, and persistently elevated IL-33 levels in mice with CEA, it is possible that IL-33 contributes to the altered neutrophil behaviour. Indeed, *ex vivo* IL-33 stimulation potentiated the phenotype in neutrophils from mice with CEA, whereas again, neutrophils from naïve mice were unresponsive. Collectively, our *in vivo* and *in vitro* findings demonstrate that neutrophils from mice with experimental asthma behave differently from those isolated from naive controls. Whether this phenotype occurs as a consequence of cytokines that ‘spill over’ from the lungs, or is mediated by factors released locally in the bone marrow, for example, IL-4 produced by resident memory Th2 cells, or a combination of both, remains to be elucidated.

The role for IL-33, neutrophils and NETs in type 2 immunity has recently gained interest in the context of parasitic helminth infections. Early in *Nippostrongylus brasiliensis* infection, a neutrophilic response is observed in the lungs as the parasite migrates through on its way to the intestine. This neutrophilic response is rapidly resolved, and an eosinophil-dominant type 2 response is evident at later time points.^32–34^ Intriguingly, this early neutrophilic response is required for effective type 2 anti-helminth immunity.^35^ Although the role of NETosis has not been investigated in the generation of type 2 immunity during helminth infection, NETosis also slows the migration of parasitic hookworms, and DNase treatment (degrading NETs) increases parasite burden.^36^ Of note, in the context of the current study, many parasites (including *H. polygyrus*) secrete DNaseII to degrade NETs: thus HpARI, HpBARI^37^ and DNaseII secreted products may all be directed against the same IL-33-NET-type-2 immunity axis.

Recent ‘omics studies have shown that neutrophils express distinct gene expression patterns depending on their developmental stage and microenvironment,^38^ and these transcriptional and epigenetic programs may support the programming of neutrophils with different effector functions. Thus, neutrophils are increasingly recognized to be more plastic and versatile than previously thought.^38–40^ For example, we previously observed that CXCR4^hi^ neutrophils, recruited to the lungs in response to low dose LPS, are the subpopulation that undergoes NETosis and promotes allergic sensitization.^41^ This raises the tantalizing possibility of targeting specific molecular pathways and programs that regulate the aberrant neutrophil response in asthma. A greater understanding of these molecular events will likely reveal novel tractable targets for therapeutic intervention. Before then, our findings expand the ‘IL-33 endotype’ in asthma to include virus-induced neutrophilic inflammation, and suggest that IL-33 blockade may be of therapeutic benefit to patients with asthma beyond the type 2-high and eosinophilic phenotypes, and potentially other diseases where neutrophilic inflammation is detrimental.

## Materials and methods

### Mouse Strains and treatments

BALB/c mice were maintained under specific pathogen free (SPF) conditions at the QIMR Berghofer Medical Research Institute Animal Facility. The mice were housed in individually ventilated cages. Pneumonia Virus of Mouse (PVM) stock J366 was prepared as previously described.^24^ Mice were inoculated with PVM (1 plaque forming unit, PFU) or vehicle diluent (10% FCS in DMEM in 10μL) via the intranasal route (i.n.) at postnatal day (PND) 7, and re-inoculated with PVM (20 pfu) at PND49 (42 days post infection, dpi) as previously described.^14^ Mice were inoculated with cockroach extract (Greer; CRE; 1μg in 50μL PBS) i.n. on 3 dpi, 45 dpi, 52 dpi, 59dpi and 66 dpi. After a rest period, the mice were then inoculated with RV1b (5×10^6^ TCID50), recombinant IL-33 (BioLegend; 40 μg/kg) or diluent (50μL PBS). In some experiments, prior to RV infection, mice were administered diluent (50μL PBS) or 160 μg/kg of HpARI protein (provided by Dr. Henry McSorley, University of Dundee, Scotland). HpARI was produced and purified as previously described.^42^ Briefly, a pSecTAG2 plasmid (ThermoFisher) was produced, encoding HpARI (Wormbase Parasite accession number HPBE_0000813301) with a C-terminal TEV site, c-myc tag and 6-His tag. The plasmid was transfected into Expi293 cells following the Expi293 expression system kit manufacturer’s instructions (ThermoFisher). Five days after transfection, supernatant was collected, and protein was purified using nickel sepharose chromatography. Purified HpARI protein was dialysed into PBS, filter sterilised and tested for endotoxin content using the HEK-Blue LPS detection kit 2 (InvivoGen) and found to have <0.002 EU endotoxin per mg of protein. To inhibit PAD-4, Cl-Amidine (Cayman Chemical; 10 mg/kg) or vehicle diluent (PBS) was administered by intraperitoneal (i.p.) injection at −24, 12, 24 and 36 hours post exposure, as described previously.^41, 43^ All experiments were approved by the QIMR Berghofer Animal Ethics Committee.

### Flow cytometry and cell sorting

The left lung lobe and the smallest postcaval lobe were mechanically dissociated using a syringe plunger and 70μm cell strainer (BD Biosciences, San Jose, CA). Isolated single cell suspensions were washed with PBS/2% FCS and red blood cells lysed using Gey’s buffer. Zombie Aqua (Biolegend, San Diego, CA) was used to exclude dead cells. The cells were then incubated with anti-FcγRIII/II (Fc block) and stained with fluorescently labelled antibodies directed against CD4-FITC (RM4.5), CD90.2-APCcy7 (53-2.1), CD200R1-APC (OX-110), NKp46-V450 (29A1.4), CD11b-PercpCy5.5 (M1/70), (all BD Bioscience, San Jose, CA), CD8-PercpCy5.5 (53-6.7), TCRγδ-APC (GL3), CD45-V421 (30-F11), TCRβ-V605 (H57-597), CD19-FITC (1D3/CD19), CD11c-FITC (N418), CD45R-FITC (RA3-6B2), Ly6G-FITC (1A8), CD11c-BV785 (N418), (all Biolegend, San Diego, CA), CD3e-FITC (145-2C11), F4/80-FITC (BM8), (all eBioscience, San Diego, CA). For intracellular staining, cells were fixed and permeabilised using the BD Cytofix/Cytoperm™kit as per the manufacturer’s instructions (BD Biosciences, San Jose, CA) and stained with GATA3-PE (TWAJ; eBioscience, San Diego, CA) and RORγT-V650 (Q31-378; BD Biosciences, San Jose, CA). Stained cells were washed and acquired on a BD LSRFortessa™(BD Biosciences, San Jose, CA) or sorted using a BD FACSAria™III (BD Biosciences, San Jose, CA). Data were analysed using FACS Diva Software v8 and FlowJo v10.8. Immune cells were identified as follows: eosinophils: CD45+, CD11b+, CD11c-, Siglec-F+, Ly6G-; neutrophils as CD45+, CD11b+, CD11c-, Siglec-F-, Ly6G+, Th2 cells as CD45+ TCRß+ CD4+ CD8-GATA3+ RORT-. Th17 cells as CD45+ TCRß+ CD4+ CD8-GATA3-RORT+ ILC2 cells as Lineage-(CD3, CD19, CD45R, CD11c, CD11b), CD45+ CD90.2+, NKp46-, GATA3+ RORT’-, ILC3 as Lineage-(CD3, CD19, CD45R, CD11c, CD11b), CD45+ CD90.2+, NKp46-, GATA3-RORT+.

### Immunohistochemistry and immunofluorescence

Paraffin-embedded lung sections were generated as previously described.^24^ For immunostaining, blocking was performed with 10% goat serum in PBS. Sections were probed with anti-Muc5ac (45M1, ThermoFisher), anti-MPO (Polyclonal, R&D Systems) or anti-CitH3 (Polyclonal, abcam) and incubated overnight at room temperature. For immunohistochemistry, the sections were washed with PBS/0.05% Tween-20 and incubated in anti-rabbit IgG-AP (Sigma-Aldrich) for 1 hour. Following incubation with appropriate secondary antibodies, immunoreactivity was developed with Fast Red (Sigma-Aldrich) and counterstained with Mayer’s hematoxylin. For immunofluorescence, the sections were washed with PBS/0.05% Tween-20 and incubated with Goat-Anti Rat IgG AF647 (ab150159, Abcam) and Rabbit Anti-Mouse IgG AF555 (A27028, Invitrogen) for 1 hour. The sections were then washed with PBS/0.5% Triton and counterstained with 4’,6-diamidino-2-phenylindole (DAPI, Sigma-Aldrich). Digitally scanned sections were analysed using Image Scope software (Scanscope XT; Aperio). Immunofluorescent slides were imaged at 20x using Zeiss 780-NLO spinning disk confocal microscope.

### Cytokine and dsDNA analysis

Bronchoalveolar lavage (BAL) fluid was collected as previously described^44^ centrifuged at 5000 rpm and the supernatant stored at −80°C prior to analysis. The concentration of IL-1α, IL-1ß, IL-4, IL-5, IL-6, IL-17A, IL-33, TSLP (Biolegend, San Diego, CA) and CXCL1 (aka KC) (R&D Systems, Minneapolis, MN) were measured in BAL by ELISA according to manufacturer’s instructions. IL-13 concentration was measured by cytokine bead array (Biolegend, San Diego, CA). dsDNA in BAL was measured by Quant-iT^™^ PicoGreen^™^ dsDNA Assay Kit (Invitrogen) according to manufacturer’s instructions.

### Neutrophil purification and culture

Blood was obtained by cardiac puncture, and bone marrow cells were harvested from the femur by flushing with a syringe. Following centrifugation (10,000 rpm for 10 mins), the cell pellet was resuspended and passed through a 70 μm cell strainer (BD Biosciences, San Jose, CA). Contaminating erythrocytes were lysed using Gey’s buffer. Neutrophils (Live, Siglec F^-^CD11b^+^Ly6G^+^CD11c^-^) were purified by FACS (>95% purity), seeded into a 12-well culture plate (50 × 10^3^/well in 1mL) in RPMI media containing 10% FCS, and cultured for 4 hours in the presence or absence of recombinant IL-33 (50 ng/ml). After centrifugation, the supernatant was harvested and stored at −80°C prior to analysis. The cell pellet was washed in PBS and fixed in 4% paraformaldehyde. Slides for immunofluorescence were prepared using a StatSpin Cytofuge as described previously.^45^

### Clinical samples

Study volunteers were recruited and inoculated with RV16 as described.^4, 11^ Nasal samples were collected together with daily respiratory symptom scores. Measurements of IL-33, dsDNA, and neutrophil elastase were performed as part of two previous studies.^4, 11^

### Statistical analysis

GraphPad Prism 8.0 software (La Jolla, California) was used for statistical analyses. Mann-Whitney test, one-way ANOVA with a Tukey post-hoc test or two-way ANOVA with Sidak post-hoc test were applied as appropriate. Correlations were evaluated using a Spearman rank correlation analysis. *p* < 0.05 was considered statistically significant.

---

**Supplementary Figure 1:**
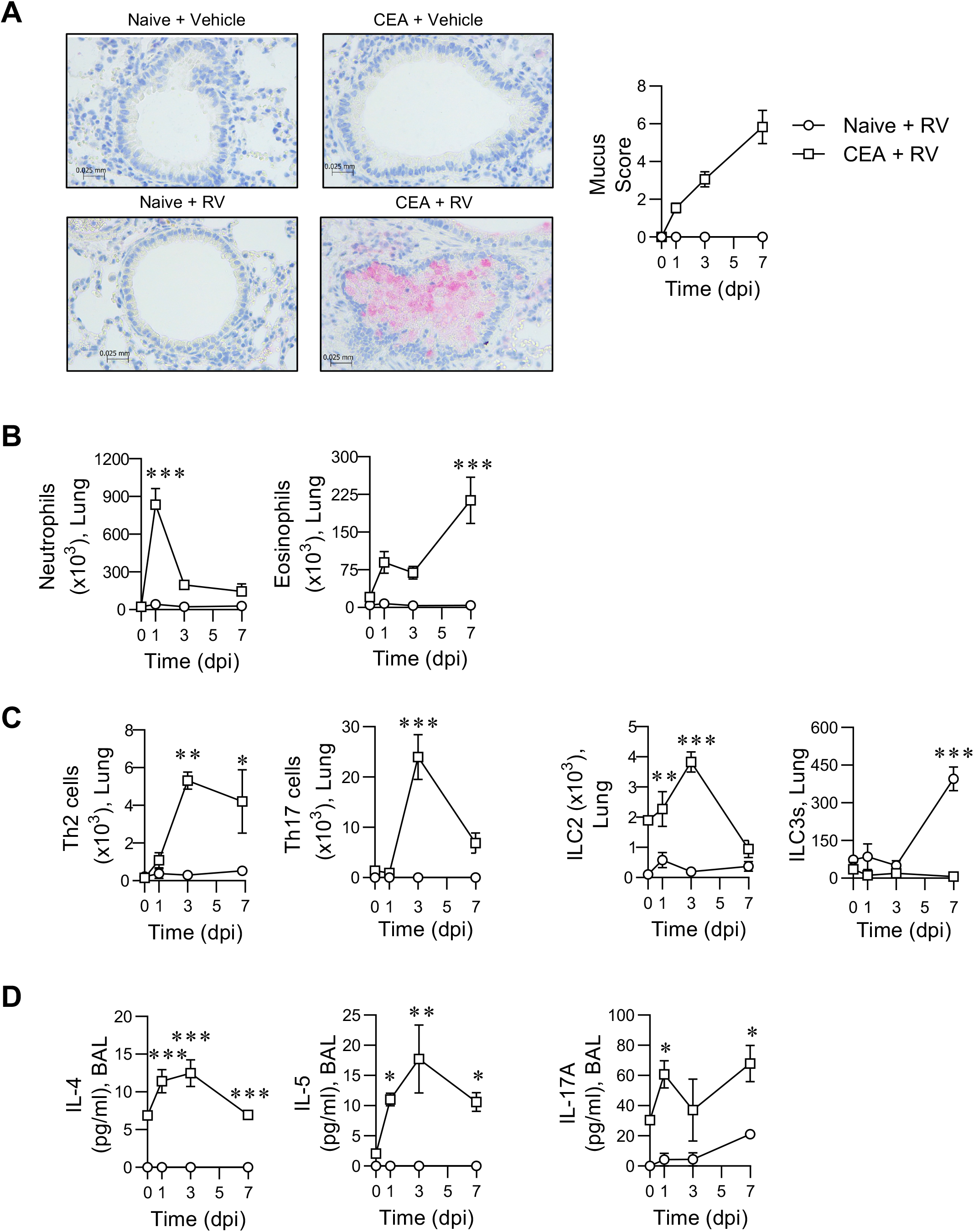
RV-induces an acute exacerbation in mice with CEA but not naïve controls. **A,** Representative images of MUC5ac immunohistochemistry (x20 magnification). **B,** Lung neutrophils and eosinophils. **C,** Lung Th2 cells, Th17 cells, ILC2s, and ILC3s. **D,** IL-4, IL-5, and IL-17A expression in BAL fluid. Data represented as mean ± SEM, *n* = 4-12 mice per group, \**P* < .05, \*\**P* < .01, *** *P* < .00.1. The data in Panel B (CEA + RV group) are the same as those presented in Fig.1C.

**Supplementary Figure 2:**
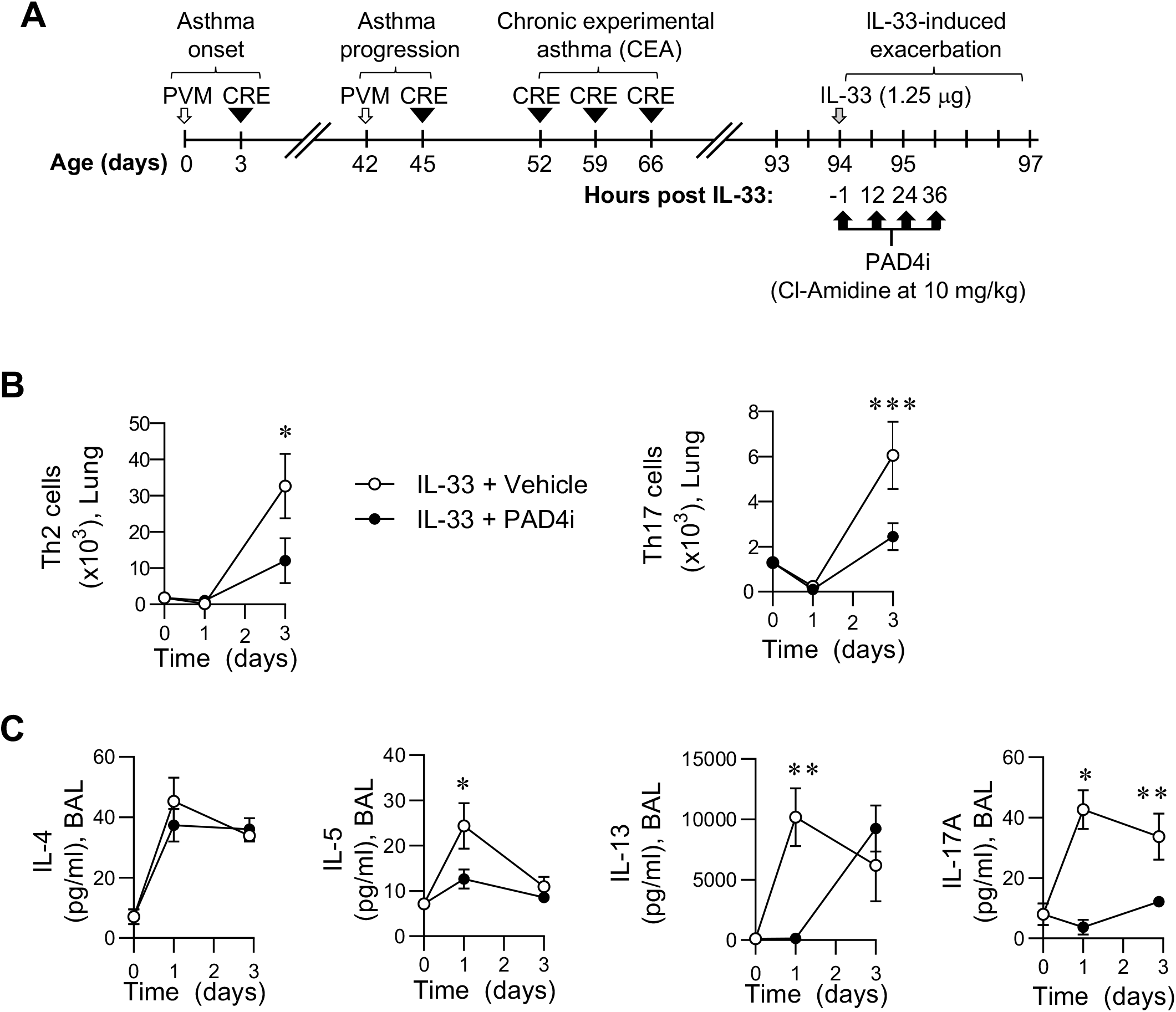
NETosis inhibition attenuates an IL-33-induced exacerbation of CEA. **A,** Study design. To inhibit NETosis mice were treated with the PAD4 inhibitor CI-Amidine (10 mg/kg) at −24, 12, 24 and 36 hours post IL-33 administration. **B,** Lung Th2 cells and Th17 cells. **C,** IL-4, IL-5, IL-13 and IL-17A expression in BAL fluid. Data represented as mean ± SEM, *n* = 4-8 mice per group, \**P* < .05, \*\**P* < .01, *** *P* < .00.1.

**Supplementary Figure 3:**
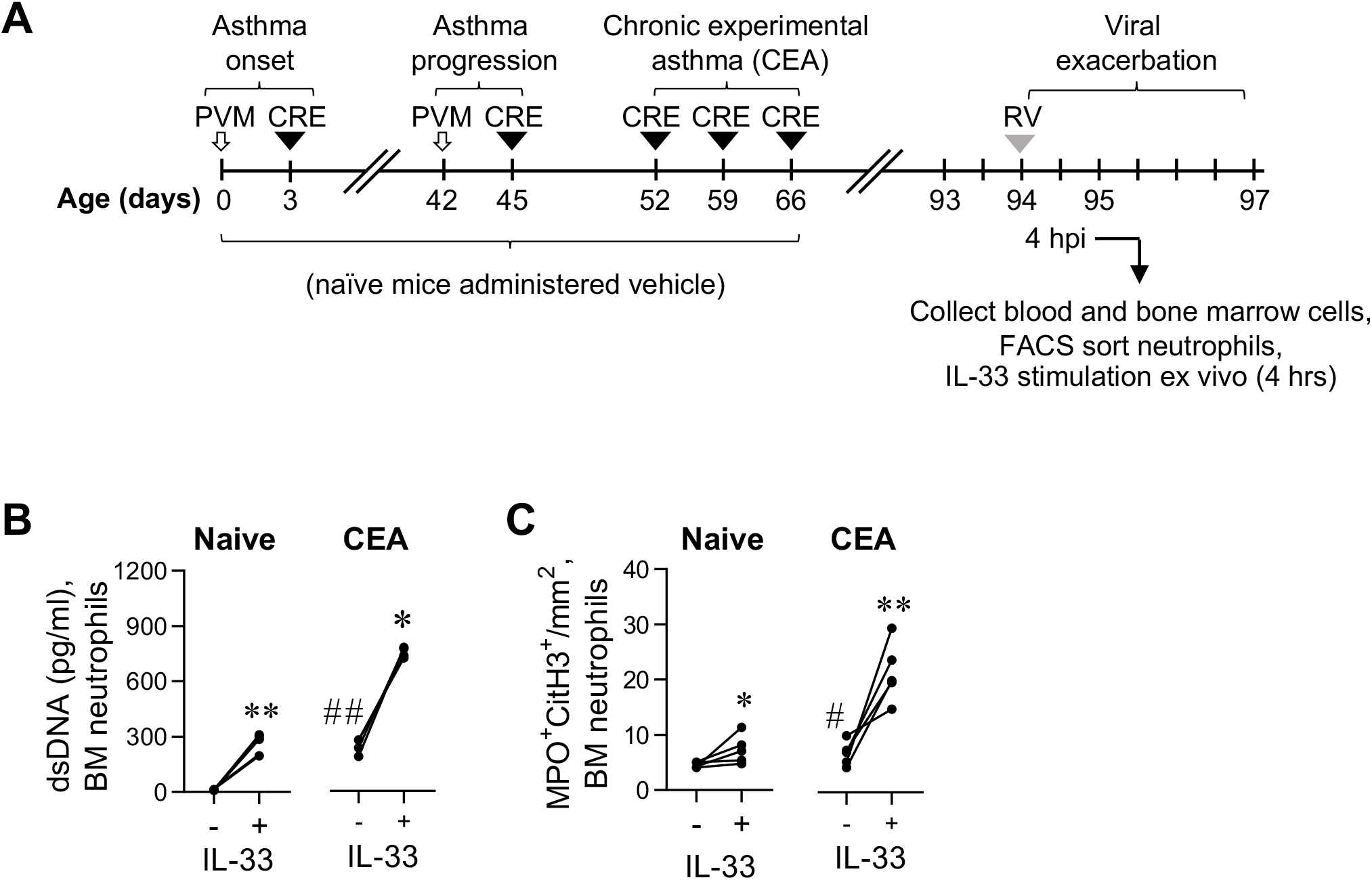
IL-33 induced NETosis of bone marrow neutrophils is greater in mice with CEA. **A,** Study design. CEA was induced through PVM and CRE exposure. At day 94, mice were inoculated with RV (5×10^6^ TCID50), and 4 hour later, neutrophils were isolated from the peripheral blood or bone marrow (BM) then stimulated with IL-33 (+) or diluent (-) for 4 hours. **B,** dsDNA levels. **C,** Quantification of co-localised MPO and CitH3 expression. Data represented as mean ± SEM, *n* = 4 mice per group, \**P* < .05, \*\**P* < .01, *** *P* < .00.1.

## Author Contributions

S.P. and H.M. conceived and designed experiments. B.C. and T.A. conducted the majority of the experiments, with support from D.H., M.A.U., I.S., R.R., M.A.A.S, A.B., and S.N. Clinical data, reagents and critical analysis was provided by M.E., S.L.J., and H.M. The manuscript was initially drafted by S.P. and B.C.; all authors contributed thereafter.

## Acknowledgments

This work was supported by an NHMRC grant awarded to S.P. (1141581), and grants to H.J.M from LONGFONDS Accelerate as part of the AWWA project, and the Medical Research Council (MR/S000593/1). The authors thank QIMR Berghofer Flow Cytometry Facility and Histology Facility for technical assistance.

---

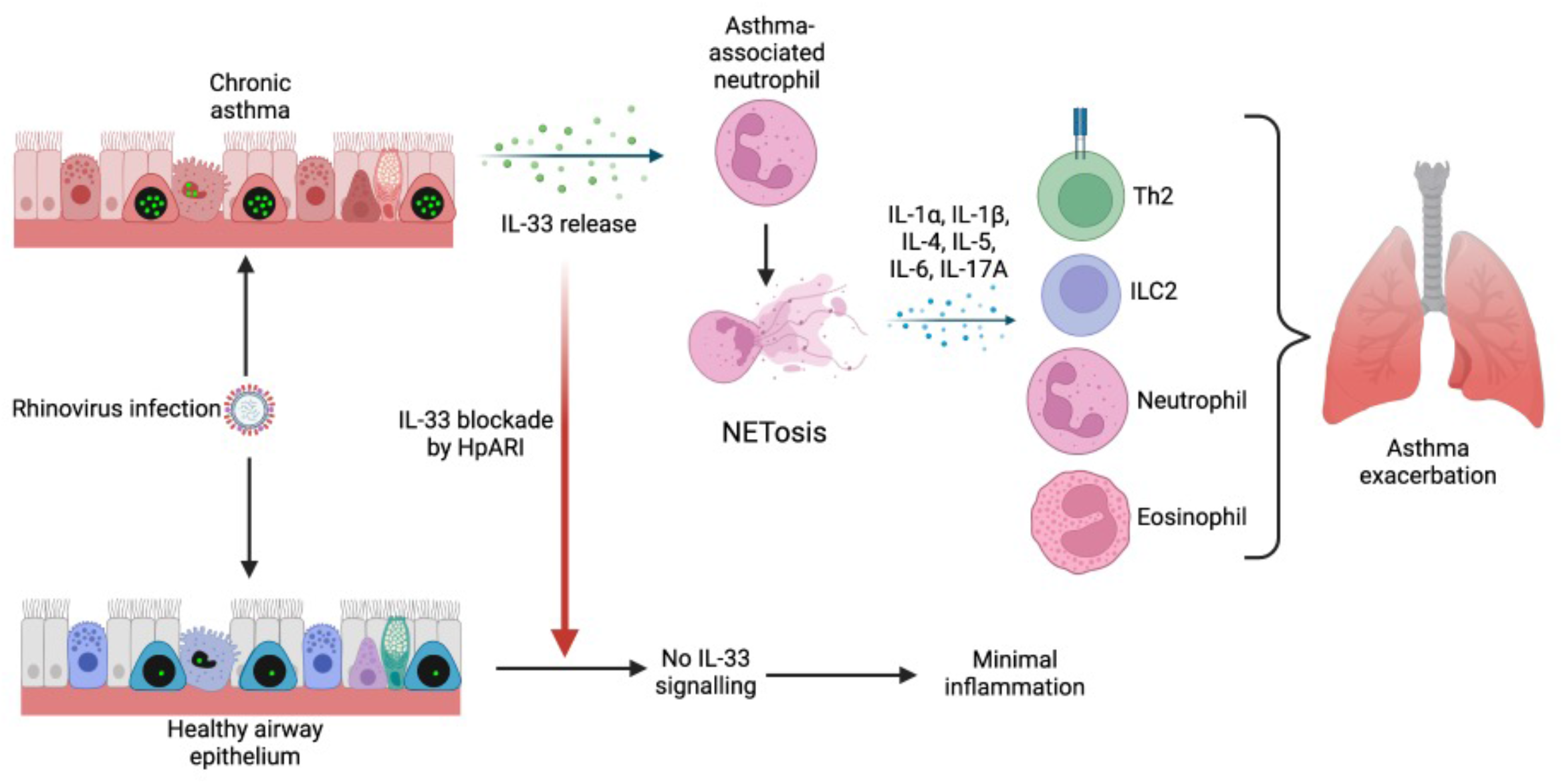
Rhinovirus (RV) infection during chronic asthma results in IL-33 release. IL-33 directly induces NETosis of neutrophils from a chronic asthma environment (but not neutrophils from a healthy subject), leading to pro-inflammatory cytokine release, neutrophilic and eosinophilic inflammation, and asthma exacerbation. By blocking IL-33, neutrophil NETosis is inhibited, and excessive inflammation is prevented.

